# SMAdd-seq: Probing chromatin accessibility with small molecule DNA intercalation and nanopore sequencing

**DOI:** 10.1101/2024.03.20.585815

**Authors:** Gali Bai, Namrita Dhillon, Colette Felton, Brett Meissner, Brandon Saint-John, Robert Shelansky, Elliot Meyerson, Eva Hrabeta-Robinson, Babak Hodjat, Hinrich Boeger, Angela N. Brooks

## Abstract

Studies of *in vivo* chromatin organization have relied on the accessibility of the underlying DNA to nucleases or methyltransferases, which is limited by their requirement for purified nuclei and enzymatic treatment. Here, we introduce a nanopore-based sequencing technique called Small-Molecule Adduct sequencing (SMAdd-seq), where we profile chromatin accessibility by treating nuclei or intact cells with a small molecule, angelicin. Angelicin reacts with thymine bases in linker DNA not bound to core nucleosomes after UV light exposure, thereby labeling accessible DNA regions. By applying SMAdd-seq in *Saccharomyces cerevisiae*, we demonstrate that angelicin-modified DNA can be detected by its distinct nanopore current signals. To systematically identify angelicin modifications and analyze chromatin structure, we developed a neural network model, NEural network for mapping MOdifications in nanopore long-reads (NEMO). NEMO accurately called expected nucleosome occupancy patterns near transcription start sites at both bulk and single-molecule levels. We observe heterogeneity in chromatin structure and identify clusters of single-molecule reads with varying configurations at specific yeast loci. Furthermore, SMAdd-seq performs equivalently on purified yeast nuclei and intact cells, indicating the promise of this method for *in vivo* chromatin labeling on long single molecules to measure native chromatin dynamics and heterogeneity.

**GRAPHICAL ABSTRACT:** 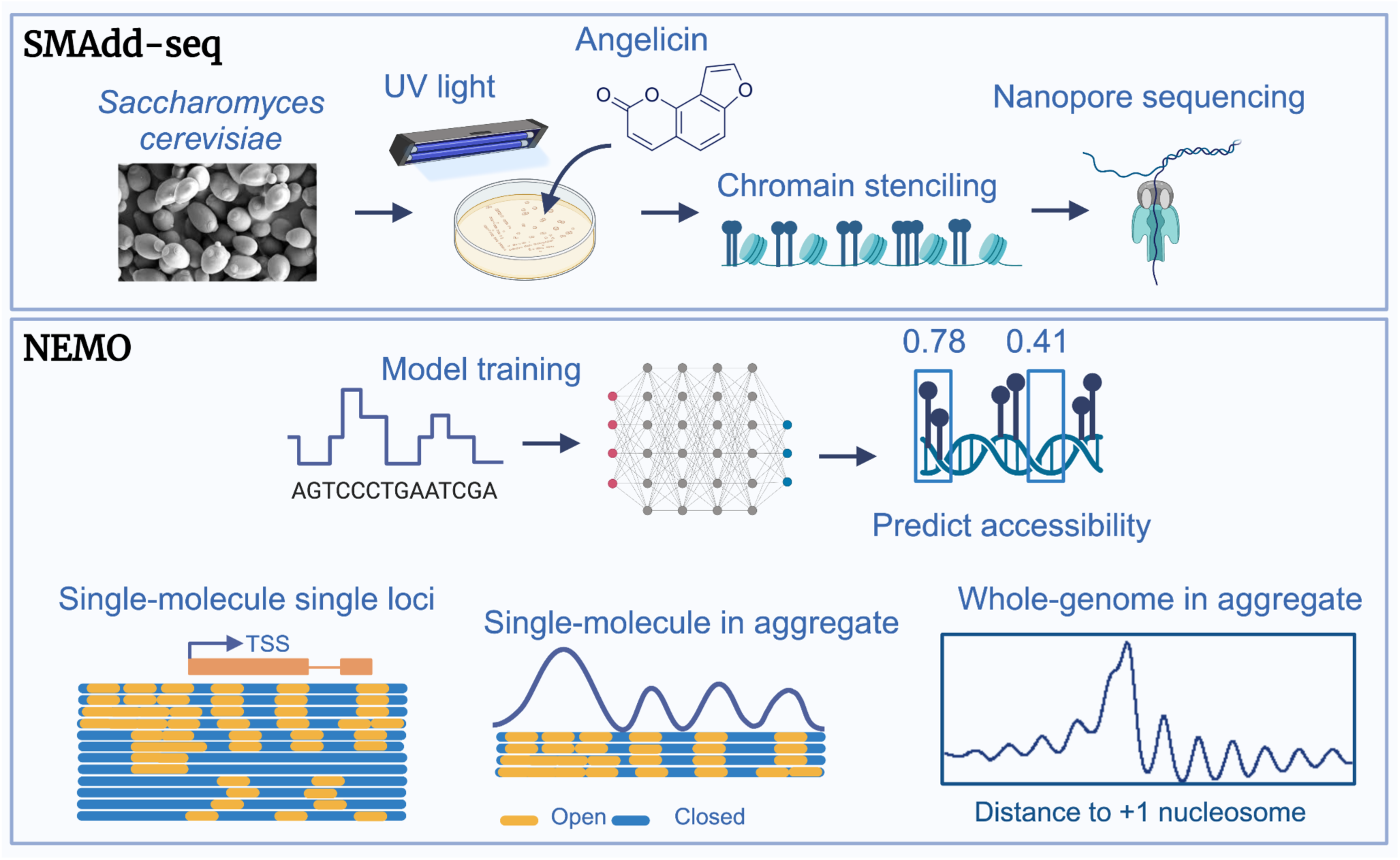

## INTRODUCTION

DNA in all eukaryotic cells is packaged into nucleosomes. This nucleoprotein complex together with DNA-binding proteins and RNA comprises chromatin. The dynamic and variable nature of chromatin regulates all DNA-centric processes and plays a vital role in cell growth, differentiation, and development. Nucleosomes are composed of approximately 147bp (∼ 1.7 turns) of DNA wrapped around a central histone protein octamer. Arrays of nucleosomes separated by ∼20-90 bp of linker DNA appear as beads on a string in electron micrographs (Olins and Olins 1974). Nucleosomes block access of DNA binding factors to the underlying DNA and impede transcription, replication, DNA repair and recombination machineries (Hughes and Rando 2014). The distribution of nucleosomes across the genome is not uniform and varies significantly between open and closed chromatin. There is also considerable heterogeneity in nucleosome distribution at different gene loci in open chromatin and also within each gene (Brown et al. 2013). This chromatin structure varies with growth conditions, differentiation, and development (Berger 2007). Thus, knowledge of the dynamic chromatin landscape can yield important insights into development, disease, and drug response.

Assays to determine nucleosome distribution at specific gene loci were developed soon after the discovery of the nucleosome (Kornberg 1974). The original assays probed for accessibility of chromatin to DNA endonucleases that mostly cleave linker DNA (C. Wu et al. 1979; Keene and Elgin 1981). These were subsequently adapted to genome-wide nucleosome distribution studies using short-read Illumina sequencing leading to MNase-seq (Johnson et al. 2006), DNAse-seq (Boyle et al. 2008), and ATAC-seq (Buenrostro et al. 2013) among others. While nucleosome distribution profiles from short-read data have been vital to our understanding of chromatin structure and function, they only provide an aggregate view of nucleosome distribution across all cells in the population. A granular view of the heterogeneity in nucleosome spacing in individual cells is lacking in these data. Also absent is a view of coordination of nucleosome organization across long genomic distances. Short-read data also suffer from biases introduced by PCR amplification, read mapping, and DNA fragmentation (Meyer and Liu 2014).

A more recent advancement in sequencing was the development of long-read nanopore sequencing technology, where an electrical current is passed across a biological pore embedded in a lipid bilayer. As single-stranded DNA is channeled through the pore by a motor protein, the electrical current undergoes shifts based on the sequence of the six bases of DNA (k-mer) present in the pore at any given time (Jain et al. 2016). Modified DNA bases can also be detected from electrical shifts from nanopore sequencing (Beaulaurier, Schadt, and Fang 2019) leading to the development of single-molecule long-read assays to map chromatin accessibility using DNA methyltransferases (MTase). Long-read sequencing approaches allow for the detection of modified DNA without the bias of PCR amplification and can also detect endogenous DNA modifications such as 6mA and 5mC (Lee et al. 2020; Stergachis et al. 2020; Yue et al. 2022; Y. Wang et al. 2019; Abdulhay et al. 2020). Data from these methods have yielded novel insights into single-cell nucleosome distribution in the genome and gene regulation.

Despite the promise of MTase assays to map nucleosome occupancy for long-read sequencing, they all rely on extracting cell nuclei before MTase treatment rendering the procedure arduous and also subject to a disrupted chromatin state (Prentice and Gurley 1983; X. Wang and Simpson 2001).

To develop a method that allows for accurate, *in vivo* chromatin labeling, we explored the use of small molecules to map accessible chromatin. The application of small molecules in chromatin structure studies dates back to the early 1900s. One such molecule, furocoumarin psoralen, intercalates between double-stranded nucleic acids and undergo photocycloaddition with thymine/uracil pyrimidine bases when exposed to UVA light to form DNA/RNA adducts (Cimino et al. 1985; Lu et al. 2016) and (Komura et al. 2001). Psoralen adducts preferentially occur in linker DNA in chromatin and this property has been widely applied in structural DNA and chromatin accessibility studies (Cimino et al. 1985). For instance, we previously used psoralen cross-linking of linker DNA in EM studies to highlight the stochastic positioning of nucleosomes on *PHO5* promoter molecules (Brown et al. 2013). Due to its extended structure, psoralen forms both covalent mono- and di-adducts with one or both strands of a nucleic acid helix with the latter forming interstrand cross-links. The psoralen family of molecules exhibits a 5’-TA > 5’-AT > 5’-TG > 5’-GT dinucleotide preference for DNA cross-links (Espisito et al 1988), with 5’-TA dinucleotides being significantly preferred.

Angelicin is an isomer of psoralen with an atomic mass of 186 g/mol (similar to glucose). Like psoralen, angelicin has been shown to preferentially form adducts with DNA not bound by nucleosomes or transcription factors (Komura et al. 2001). Unlike psoralen, angelicin, due to its angular structure, is described to form photo-monoadducts with one of the two DNA strands (Komura et al. 2001; Ashwood-Smith and Grant 1977). While some studies report that angelicin can slowly form covalent interstrand cross-links upon prolonged UV irradiation (1 hour), the authors recommend its application for photoinduced-labeling of chromatin in instances where covalently cross-linked DNA strands are problematic (Lown and Sim 1978). Furthermore, angelicin easily traverses cell membranes and thus can be applied to analysis of chromatin structure with little perturbation of the cells (Cimino et al. 1985) (Cleaver 1985) and (Cleaver 1985; Komura et al. 2001). We exploited the cell permeability of angelicin as well as its ability to primarily form photoadducts on single strands of accessible DNA to assess whether we can detect these adducts using nanopore sequencing.

Here we report the development of SMAdd-seq that utilizes angelicin to map chromatin accessibility with nanopore sequencing. We show that intercalation of angelicin causes a detectable shift in the nanopore current signal and have developed a neural network approach to predict angelicin modification from these signal data. We can accurately detect known chromatin accessibility patterns in angelicin-modified nuclei and intact cells (spheroplast) in *S. cerevisiae* and can discern single-molecule chromatin profiles and regulatory patterns at individual loci.

## MATERIAL AND METHODS

### Yeast strains and culture

The *Saccharomyces cerevisiae* strain YS18 (MATalpha his3-11 his3-15 leu2-3 leu2-112 can1-100 ura3Δ5) (S288C derivative) was used in this study. Cells were grown in YPD (1% yeast extract, 2% peptone, 2% dextrose) at 30°C.

### Yeast spheroplast preparation and nuclei isolation

Yeast spheroplast preparation and nuclei isolation were carried out as described previously (Brahma and Henikoff 2022).

### Angelicin modification of yeast and genomic DNA extraction

Yeast chromatin was modified with angelicin using either purified nuclei or yeast spheroplasts. For yeast spheroplasts, fresh spheroplast yeast from a 250-mL culture were resuspended in 0.4 mL of angelicin modification buffer (10mM Tris-HCl, 10mM NaCl, 0.1mM EDTA, pH 7.4). The spheroplast suspension was divided into 3 wells (200ul each) of a 6-well cell culture plate and placed on ice. 10 uL of a 2mg/ml angelicin stock (500uM) (SIGMA-A0956) were added to each of the wells and the plate was subject to UV treatment 7 times; each separated by a 5-minute incubation on ice. Isolated Yeast nuclei were instead treated with angelicin in 7 rounds up to 1.28mM; 2 aliquots of nuclei (each containing ∼5 X 10^8 nuclei) were pooled then pelleted before resuspending in 600 uL of angelicin modification buffer. The suspension was split equally into 3 wells of a 12-well culture plate where angelicin was then added 20uM at a time to each well on the culture plate between 7 rounds of UV-cross-linking with 5-minute incubations on ice between rounds of cross-linking. Angelicin is photolytic, and care was taken to ensure samples incubating with angelicin were kept away from direct UV light before and after cross-linking. The plate was swirled a few times to mix the angelicin and incubated in the dark on ice for 5 minutes. While ensuring the culture plate remained nested in ice, the plate was exposed to 365nm UVA light (Stratagene UV Stratalinker 2400, power 5.0) for 5 minutes followed by a 5-minute incubation on ice. Both, angelicin and UV treatment steps or UV treatment alone were repeated for a total of 7 times. Contents of all wells were pooled into a fresh low-adhesion tube (EPPENDORF-022431021) and both wells were washed with 100 uL of ice-cold angelicin modification buffer and added to the same low-adhesion tube to maximize nuclei retrieval. Nuclei treatment with 500uM of angelicin was performed in biological triplicate. High molecular weight DNA was purified using the NEB Monarch HMW DNA Extraction kit for tissue (T3060L). The use of wide-bore pipette tips when working directly with long DNA significantly improves the length of the purified library. Purified DNA was quantified using the ds DNA BR kit for Qubit (ThermoFisher) and analyzed on a genomic DNA ScreenTape on a TapeStation 4150. Positive and negative control data for the neural network training were generated from purified high molecular weight DNA (∼6ug) that was incubated with or without (mock treated with ethanol) 500uM angelicin, respectively, followed by UV treatment as described above.

### Oxford nanopore sequencing

3-4 micrograms of high molecular weight DNA were used to prepare genomic libraries for sequencing with Oxford Nanopore Technologies (ONT) SQK-LSK110 kits for use with R9.4.1 (FLO-MIN106) flowcells. ∼1.5 micrograms of the library were loaded onto flowcells, and all library sequencing was undertaken on a MinION for 24 hours each with MUX scanning every 6 hours to extend the life of the flow cell.

### Basecalling and aligning sequencing data

The data was basecalled and aligned to sacCer3 genome with Dorado v0.6.2 using pore model dna_r9.4.1_e8_sup@v3.6 and parameters--emit-moves (Li 2018, 2021). Reads were aligned, sorted and indexed using Samtools v1.13. Secondary and supplemental reads were further filtered. Uncalled4 v4.1.0 was run to align signals to k-mers and obtain eventalign output (Loman, Quick, and Simpson 2015; Kovaka et al. 2025).

### Alkaline agarose gel electrophoresis

Agarose gel electrophoresis was performed on plasmid pBlueScript DNA modified with either 0, 100, 200, 500, and 1000 uM angelicin and digested with Not1 according to Green and Sambrook, 2021. The modified DNA was purified on SPRI beads and resolved on a 1% alkaline agarose gel. DNA was visualized and documented on a BioRad ChemiDoc XRS imager after ethidium bromide staining and destaining.

### Identification of k-mer signal distribution peaks and informative k-mers

We aggregated the mean signal value for each k-mer (6-mer) in each read from the eventalign file. For each k-mer, the signal density was calculated using numpy np.histogram function at the range of 25 to 150. We then used scipy.signal.find_peaks with parameters prominence=0.005 and distance=5 to identify k-mers with a secondary peak in the positive control sample. We considered these k-mers informative k-mers to indicate signal shifts produced by modification. We also identified peaks in the negative control sample and found that no k-mers had more than one peak in that sample. We then generate sequence logos using all 169 informative k-mers (X. Wu and Bartel 2017).

### Picoamp signal preprocessing for model training

Signal picoamps with a value smaller than -50 or larger than 150 were clipped to -50 and 150. Signals were normalized by the mean and standard deviation within the data input to the model. In each read from each sample, picoamp signals were scanned by a sliding window of 400 with a step size of 1. Input signals of length 400 are represented as a one-dimensional array [1, 2, 3, 4, 5, …, 400]. For every data point, a single signal shift was applied to capture the sequential nature of nanopore signals (e.g., [2, 3, 4, 5, 6, …, 401]).

### Training, validation, and testing of the neural network model

We developed NEMO (a NEural network model for mapping MOdifications in nanopore long-reads), a computational tool for training and predicting angelicin modification sites. NEMO is implemented in PyTorch (v2.0.1) (Paszke et al. 2019) and utilizes a Residual Network classifier optimized for one-dimensional signal data analysis (He et al. 2016; Hong et al. 2020). Each positive and negative control dataset was divided into train (60%), validation (20%) and test (20%) sets. Positive control data were labeled with prediction probabilities of 1.0, and negative control data were labeled with prediction probabilities of 0.0. The model was trained over 100 epochs, with a batch size of 512 and 1000 batches per epoch. For training model parameters with gradient descent, binary cross-entropy loss was calculated using the function torch.nn.BCELoss after each step, and model parameters were updated with the function torch.optim.Adam (Kingma and Ba 2014). Following each epoch of training, model performance was evaluated in the validation set with batch size of 512 and 500 batches per epoch. Training accuracy, training loss, validation accuracy and validation loss were recorded per epoch. The model with the highest validation accuracy after 100 epochs was saved as the optimal model for further analyses.

This final model was applied to classify signals in the test dataset, where standard performance metrics including True Positive Rate (TPR), False Positive Rate (FPR), True Negative Rate (TNR), False Negative Rate (FNR), accuracy, precision, recall, and F1 score were calculated to comprehensively assess the model’s performance. ROC and AUC scores were generated using the roc_curve and auc functions in scikit-learn (Pedregosa et al. 2012).

### Neural network prediction in chromatin sequencing data

The model trained on control datasets was applied to predict angelicin modifications in normalized chromatin sequencing data. First, for each signal sequence mapped to a single read, NEMO scans the signals with a sliding window of 400, moving in steps of 200. Then, each length-400 signal was fed into the neural network model to receive a modification probability score, which is assigned to all genomic positions within that window based on Dorado basecaller’s sequence-to-signal mapping (approximately 30 ∼ 50 bp regions). Finally, read modification scores assigned to the same genomic position are averaged to produce the final modification score for that position.

### Aggregate analysis of +1 nucleosomes dyad

For aggregate analyses, we used all annotated +1 nucleosome dyads across the yeast genome (Chereji et al. 2018). For each gene, we calculated the average modification probability for positions up to 2000 bp upstream and downstream relative to each +1 nucleosome. Finally, the scores from an individual gene locus were averaged to generate genome-wide aggregated modification scores at +1 nucleosome dyads. PacBio methylation data were downloaded from GSE243114 (Dennis, Xu, and Clark 2024). The bam files were parsed by pysam (Li 2009, Bonfield 2021 and Danecek 2021) to obtain base-level modifications, which were further averaged for positions up to 2000 bp upstream and downstream relative to each +1 nucleosome. For visualization, all averaged scores were further normalized by mean and standard deviation.

### Quality control (QC) of chromatin sequencing data

We used the observance of expected nucleosome periodicity as QC for the ability of the trained model to predict chromatin accessibility in each chromatin-modified sample. From three replicates of angelicin-treated nuclei, nucleosome occupancy patterns in replicate 3 showed poor Pearson correlation with the expected nucleosome periodicity, thus did not pass our QC (Supplemental Table 2).

### Identification of genes with well-positioned nucleosomes at transcription start sites (TSS)

To identify genes with well-positioned nucleosomes at their promoters, we calculated the Pearson correlation coefficient between each gene’s individual angelicin modification scores and the genome-wide aggregated scores within 600 bp upstream and downstream of the +1 nucleosome dyads. We also characterized the heterogeneity of nucleosome positioning at gene TSS by calculating the variance of angelicin modification scores between reads for each gene (Supplemental Table 2). Genes were first ranked by the highest Pearson correlation to the lowest, then by the height read coverage to the lowest, and by least heterogeneity to the largest. We used the 75% percentile cutoff for each sample to select genes with high Pearson correlation, high coverage, and low variability. We identified 380 genes in replicate 1, and 230 genes in replicate 2. Among those, 38 genes were shared by two replicates and used for visualization.

### Single-molecule clustering and visualization

In NEMO, we implemented a genome track visualizer using matplotlib v3.6.2 (Hunter, 2007) to show modifications in individual reads. Reads covering a minimum of 80% of given genome regions were used to construct a modification probability matrix. Highly modified reads were identified as reads with an average modification score of 0.8 and were filtered out. Missing values in the matrix were imputed with scikit-learn v1.1.2 simpleImputer function under ‘most_frequent’ strategy. The matrix was then input to the scikit-learn K-Means clustering algorithm, where reads are clustered based on their modification profiles. We initially set k = 2 for clustering reads per gene and increased k if visual inspection revealed distinct subclusters. Clustering was performed with random centroid initializations, and the cluster ids are collected after 300 iterations. Single molecules were colored based on their predicted angelicin modification scores ranging between a probability of 0 to 1, with dark blue indicating 0% probability of angelicin modifications, thus nucleosome occupied regions, while bright yellow indicating a 100% probability of angelicin modifications and thus accessible regions.

To visualize 38 genes with well-positioned nucleosomes at TSS, reads mapped to each gene promoter region (1200bp centered at +1 nucleosome dyads) were stacked together. Genes were ordered from highest to lowest Pearson correlation coefficient. Reads within each gene were ordered by k-means clustering. Aggregated angelicin modifications were calculated by averaging scores across reads for each gene, and then averaging across the genes. The averaged scores were further normalized by min-max value.

To visualize individual gene locus reads mapped to *zz-YIL161W* (chrIX:38868-40068), *FEX2* (chrXVI:13765-14965)*, CLN2* (chrXVI:66400-67550), and *NUP170* (chrII:74300-75800) are clustered and visualized using NEMO with the method described above. Aggregated angelicin modifications for each cluster were calculated by averaging scores across the reads for each cluster and plotted on top of each cluster. The averaged scores were further normalized by min-max value.

## RESULTS

### Angelicin modification and sequencing of DNA

To determine whether angelicin-modified bases can be detected through nanopore sequencing, we first prepared naked DNA treated with angelicin and 5 minutes of UV exposure for seven consecutive rounds. High molecular weight genomic DNA purified from yeast was treated with either 0uM, 20uM, 100uM, or 500uM angelicin and exposed to UV-A at 365nm for photocycloaddition (Figure 1A, Supplemental Figure 1A, Supplemental Table 1). The parameters used for angelicin treatment were those previously optimized (Brown et al. 2013) for mapping nucleosomes by *in vitro* psoralen cross-linking to ensure high levels of covalent modification of the DNA (Supplemental Figure 1B). These samples were then used to prepare libraries for nanopore sequencing using the DNA ligation kit SQK-LSK110 kit (Oxford Nanopore Technologies) and were sequenced on R9.4.1 Minion flow cells. All reads were basecalled and mapped to the yeast SacCer3.0 genome with Dorado 0.6.2 (The Nanopore Sequencing Group 2023) and then aligned to the yeast SacCer3.0 genome with Uncalled4 eventalign (Kovaka et al. 2024) (Figure 1A).

**Figure 1.**
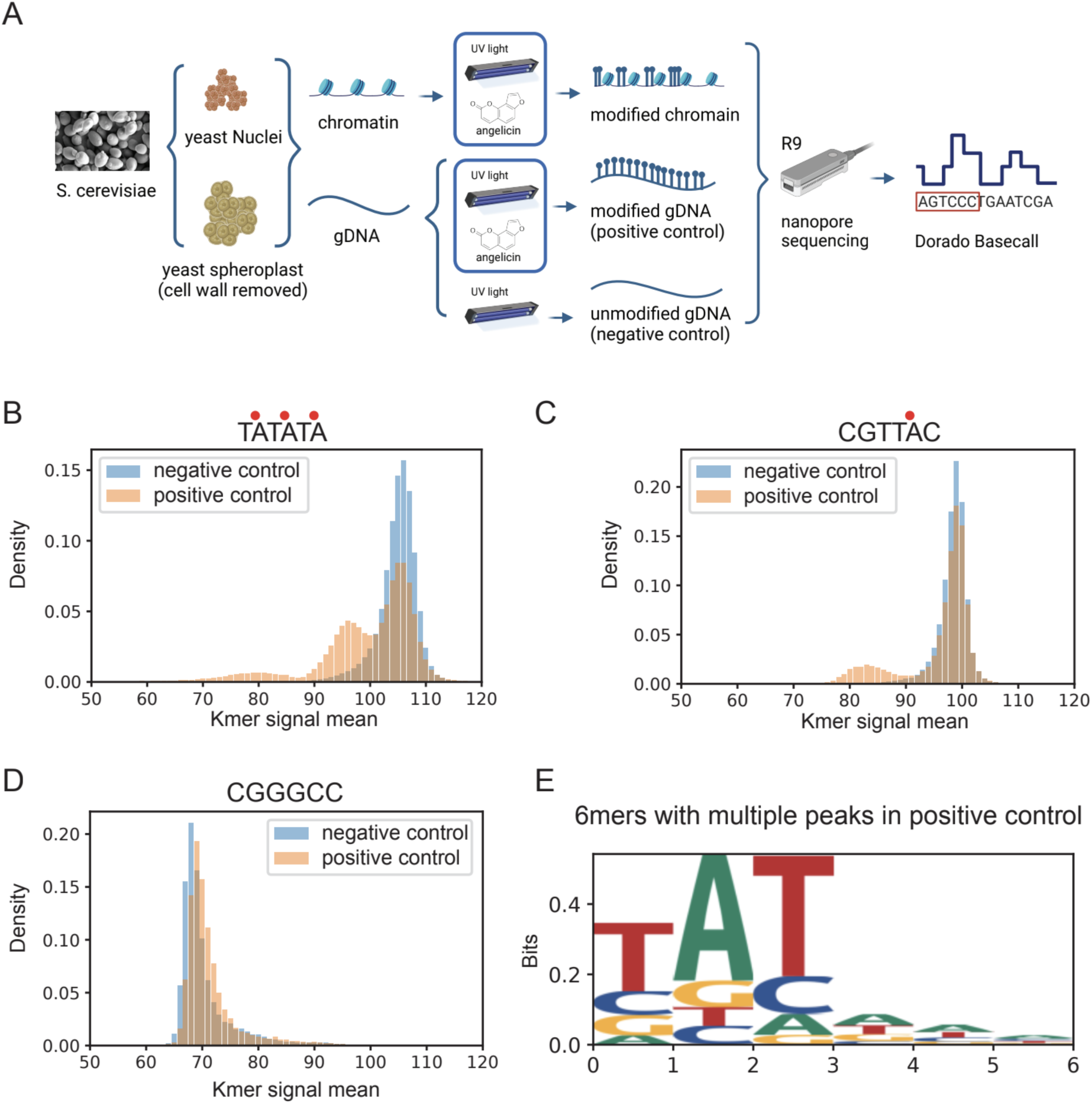
SMAdd-seq: A method using angelicin modification to probe chromatin accessibility. (A) Schematic of the SMAdd-seq method. Yeast nuclei and spheroplasts were treated with 500uM angelicin, then exposed to multiple rounds of UV light to crosslink the angelicin with DNA. The modified DNA was extracted and sequenced by nanopore sequencing with R9 flow cells, then base called with Dorado. Yeast genomic DNA were isolated and treated with 500uM angelicin under UV light or UV light only to produce positive and negative control samples. (B, C & D) Histograms of the nanopore signal currents produced from a given k-mer in yeast DNA that had been either treated with UV light only (blue: negative control) or with angelicin and UV for (orange: positive control). Red dots indicate the inferred angelicin cross-linking sites. (B) A modifiable k-mer TATATA has two shifted peaks in the positive control sample, (C) a modifiable k-mer CGTTAC has one shifted peak in the positive control sample, and (D) an unmodifiable k-mer CGGGCC has no shifted peak in the positive control sample. (E) Sequence logo for the 169 k-mers with shifted peaks in the positive control sample.

While we were able to sequence angelicin-modified DNA through the nanopore, we noticed that the sequencing output of angelicin-modified DNA was lower than unmodified DNA and the pores became inactive considerably faster (Supplemental Figure 2A-C). Given that our preliminary structural analysis showed thymine bases modified by angelicin could fit through a nanopore (data not shown), as well as the fact that we were able to sequence some angelicin-modified DNA, it was unlikely that the angelicin monoadduct was blocking the pores. Although previous work has shown that angelicin should form covalent bonds with a single thymine base on one DNA strand, without forming cross-links between the two strands of DNA (Ashwood-Smith and Grant 1977; Komura et al. 2001), a study from Lown and Sim showed that a small population of cross-linked molecules are also generated by angelicin albeit in much lower proportion compared to psoralen (Lown and Sim 1978). To assess whether angelicin induces DNA cross-linking, we asked whether we could visualize cross-links in angelicin-modified plasmid DNA on a denaturing alkaline agarose gel. This technique denatures DNA into single strands and any DNA molecules with cross-links are unable to separate into single strands and thus run slower than their single-stranded counterparts. (Supplemental Figure 2D). We treated plasmid DNA with varying concentrations of angelicin (0uM to 1mM) followed by UV exposure. While a majority of the angelicin-modified plasmid DNA molecules migrated as single strands, we did observe a small fraction of the DNA that migrated much slower than the single-stranded species suggestive of interstrand cross-links in concordance with the observation made by Lown and Sim (1978). We hypothesize that this small fraction of cross-links was sufficient to cause the rapid decay of nanopores. However, despite this reduced throughput, we were able to sequence and align about 64K reads with an average coverage of 13 from the positive control sample modified with 500uM angelicin (Supplemental Figure 1C, Supplemental Table 1).

### Identification of angelicin modification from the nanopore current signal

Using the aligned current signal data from the 500 uM positive and negative control samples, we compared the distribution of current signal values for all 6 base-pair long k-mers including those with the intercalation motif for angelicin (5’-TA, 5’-AT, 5’-TG and 5’-GT). A subset of k-mers containing the 5’-TA showed a secondary peak in the positive control due to changes in current signals (Figure 1B-C). In contrast, k-mers without the intercalation motif showed no detectable changes in current signals between the positive and negative controls (Figure 1D, Supplemental Figure 1D-E). We were somewhat surprised that we only observed a shift in 5’-TA k-mers and no discernible shifts in current with 5’-AT, 5’-TG or 5’-GT containing k-mers that should also be potentially modifiable. However, psoralens have also been shown to preferentially cross-link 5’-TA dinucleotides significantly over the rest (Esposito 1988, Komura, 2001). It is possible that non-TA k-mers were not being modified in our experimental settings or the current shifts are too subtle to be detected.

However, since we observe a current signal shift in k-mers with the 5’-TA dinucleotide and given previous work showing that other base modifications e.g. methylation, alter the current signal (Lee et al. 2020; Y. Wang et al. 2019), we concluded that this secondary distribution was due to angelicin-modified DNA. We observed a subset of k-mers with multiple 5’-TA motifs that showed three total peaks, suggesting both single and double modification by angelicin on these k-mers (Figure 1B). However, a majority of TA-containing k-mers did not show any shifts in the signal distribution between the negative and positive control (Supplemental Figure 1F).

After generating a sequence logo for k-mers with multiple signal peaks, we found the 5’-TA motif as expected, but more surprisingly, we found that the detection was limited primarily to the more specific motif 5’-TAT, which may explain why other TA-containing k-mers did not show multiple signal peaks after intercalation of angelicin (Figure 1E). From the 20uM, 100uM, and 500uM titrations of angelicin treatment on purified genomic DNA, we found that the number of informative kmers directly correlated with the amount of angelicin the DNA was treated with (Supplemental Figure 1G). However, at higher angelicin concentrations, we also had a significantly faster loss of nanopores (Supplemental Figure 2D). Therefore, we did not treat or sequence DNA with angelicin concentrations greater than 500uM.

### Identification of angelicin modification using a neural network model

Due to only being able to detect distinct angelicin-modified signal distributions for 14% of TA-containing k-mers and 4% of all k-mers based on signal density (Supplemental Figure 1F), we hypothesized that angelicin modification could more easily be detected using a machine learning model to observe changes in nanopore current signal across a larger window of bases. To map single-molecule chromatin accessibility at nucleosome resolution, we trained and tested a neural network model we call NEMO, to predict angelicin modifications from signal picoamp data without the information of underlying bases. It then maps the modification scores from signal space to sequence space based on the Dorado move-out table.

In the hyper-parameter tuning step, we tested three different model architectures with input signal lengths of 400 or 100. Since we were interested in identifying the presence or absence of nucleosomes, we picked a 400 signal measurement window as the maximum input window, which corresponds to approximately 75 bp, or half of a nucleosome footprint (Supplemental Figure 3A). The one-dimensional residual neural network (ResNet1D) (Hong et al. 2020) with an input signal length of 400 outperformed the others based on validation accuracy (Supplemental Figure 3B-C). ResNet1D has been used to monitor electrocardiogram (ECG) signal data in intensive care units (Hong et al. 2020). Considering the analogous nature of electrical current measured by electrocardiograms and ONT flow cells, ResNet1D is ideal for learning signal changes caused by nucleic acid modifications. To infer angelicin-modified regions, we trained the ResNet1D model directly from windows of consecutive nanopore signals (Figure 2A). Our positive and negative control data were used to train, validate, and test the classification ability of the neural network model. Our model was able to distinguish signal currents from positive and negative control data with an area under the receiver operating curve (AUC) of 0.9 in the stand-alone test dataset (Figure 2B). This represents a relatively high true positive rate and low false positive rate for detecting angelicin modification sites (Supplemental Figure 3D).

**Figure 2.**
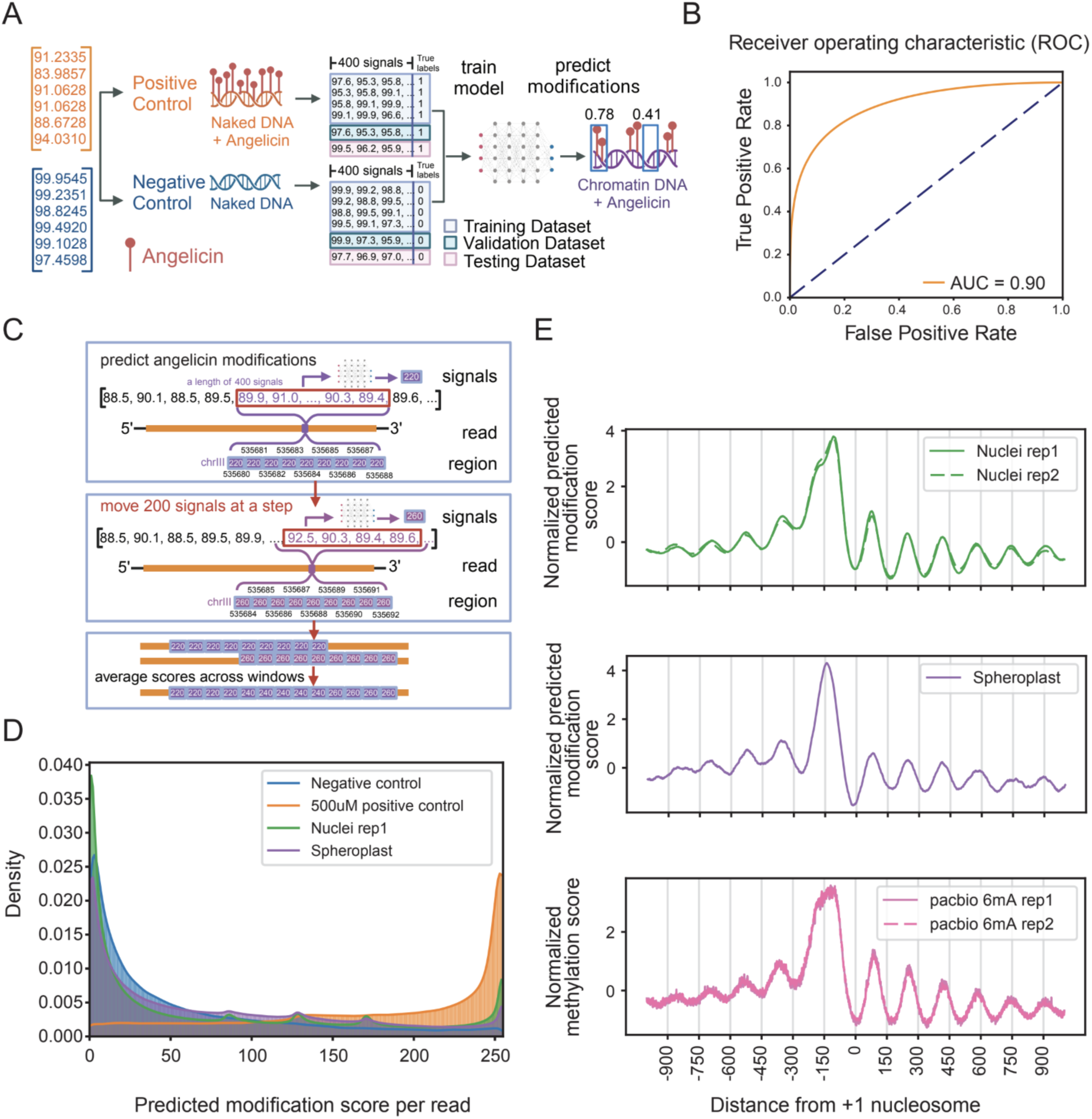
Angelicin modification scoring from the neural network model identifies expected patterns of nucleosome occupancy near transcription start sites. (A) A schematic of the neural network model trained on the nanopore signal currents produced from positive and negative control DNA sequencing. (B) Receiver operating characteristic (ROC) curve showing the performance of the trained model on the standalone test dataset. (C) A schematic showing how modification probability is predicted from a window of 400 signals per read. The scores were mapped to the genome reference and averaged per position for each read. (D) Density of average predicted scores per read for negative control DNA, positive control DNA, nuclei chromatin replicate 1 and spheroplast chromatin data. (E) Aggregate modification probability predicted by NEMO on angelicin-modified chromatin (top and middle) and PacBio 6mA methylated chromatin (bottom, data from *Clark 2024*) for 2000 base pairs centered on +1 nucleosome dyad.

### Identification of angelicin-modified DNA from nuclei and intact cells using NEMO

To evaluate the use of angelicin intercalation to determine chromatin structure, we treated purified yeast nuclei with 500uM angelicin and 5 minutes of UV exposure for seven consecutive rounds. We also treated yeast spheroplasts with 500 uM angelicin and 7 consecutive rounds of UV exposure to determine the feasibility of angelicin adduct formation in intact cells (Figure 1A). Following angelicin treatment of yeast chromatin in nuclei or spheroplasts, high-molecular-weight DNA with a mean length of ∼40kbp was extracted (Supplemental Figure 1B). DNA libraries were constructed, sequenced, and base-called the same way as the control samples. Overall, the median read length was ∼3kbp, with a median of 100k aligned reads (primary alignments) and a median base phred quality score of 22 (range from 0 to 40) (Supplemental Figure 1C, Supplemental Table 1).

We next applied NEMO to predict accessible regions in angelicin-modified chromatin data. Individual reads were scanned using a 400 signal sliding window with a 200 signal step size. Prediction scores were assigned to every base represented by the signal window, and then scores across multiple windows for each base were averaged (Figure 2C). This corresponds to an average read-level resolution of about 20bp. After applying the model to chromatin from both nuclei and spheroplasts, we saw that both showed a pattern of predicted accessibility that fell between the positive and negative control datasets, indicating an intermediate pattern of modification as expected with some genomic regions blocked from intercalation by nucleosomes (Figure 2D). The fact that the spheroplast data showed any predicted modification indicates that angelicin is, in fact, cell-permeable and could successfully modify chromatin *in vivo*.

### Identification of chromatin structure using angelicin modification

The region around the transcription start site (TSS) of a transcriptionally active gene shows a characteristic pattern of chromatin accessibility upstream of the TSS. The DNA is rendered generally accessible, allowing general transcription factors and RNA polymerase II to bind. Downstream of the TSS is the +1 nucleosome, followed by a regular pattern of positioned nucleosomes interspersed with accessible linker regions. Furthermore, the first nucleosome is expected to be the most well-positioned, with subsequent downstream nucleosomes being less well-positioned (Chereji, Ocampo, and Clark 2017). After generating accessibility predictions for each read from the neural network, we averaged all predictions for a window of +/-1000 bp around each +1 nucleosome at protein-coding genes and normalized its mean and standard deviation (Figure 2E). From this metagene plot, we found that both the angelicin-modified nuclei and spheroplast chromatin samples closely replicated the pattern of accessibility expected from previous short-read methods and with a recently published orthologous long-read method using EcoGII to detect chromatin accessibility with PacBio sequencing (Dennis, Xu, and Clark 2024) (Figure 2E). We also found that this method is reproducible, giving a similar pattern between our nuclei sequencing replicates (Figure 2E). While investigating the cause of a small dip at -150bp in the metagene plot from nuclei, we found that yeast TSSs are mostly enriched for the 5’-TA motif except at position -150bp upstream of the +1 nucleosome, which would bias the angelicin modification efficiency (Supplemental Figure 3E). Curiously, we did not observe this dip in our spheroplast sample and future investigations will explore whether this is a technical or biological difference, given that we are modifying chromatin, *in vivo*, with spheroplasts.

### Identification of heterogeneous chromatin accessibility patterns at individual genes

To examine patterns of accessibility on a single-gene, single-molecule level, we examined our nuclei replicate data, which had the highest read coverage. We identified genes with well-positioned +1 nucleosomes by identifying the Pearson correlation of the accessibility pattern at the gene level with the accessibility pattern at the metagene level (Supplemental Figure 4, Supplemental Table 2). We identified 38 genes with significant correlation of their accessibility pattern in both nuclei replicates (replicate 1 and 2) and visualized the window around the +1 nucleosome for every read aligned to these genes (Figure 3A). We show that the angelicin modification-based accessibility predictions represent the expected chromatin structure even on the single-read level. The promoter and +1 and +2 nucleosomes can clearly be seen from the read accessibility across these top genes.

**Figure 3.**
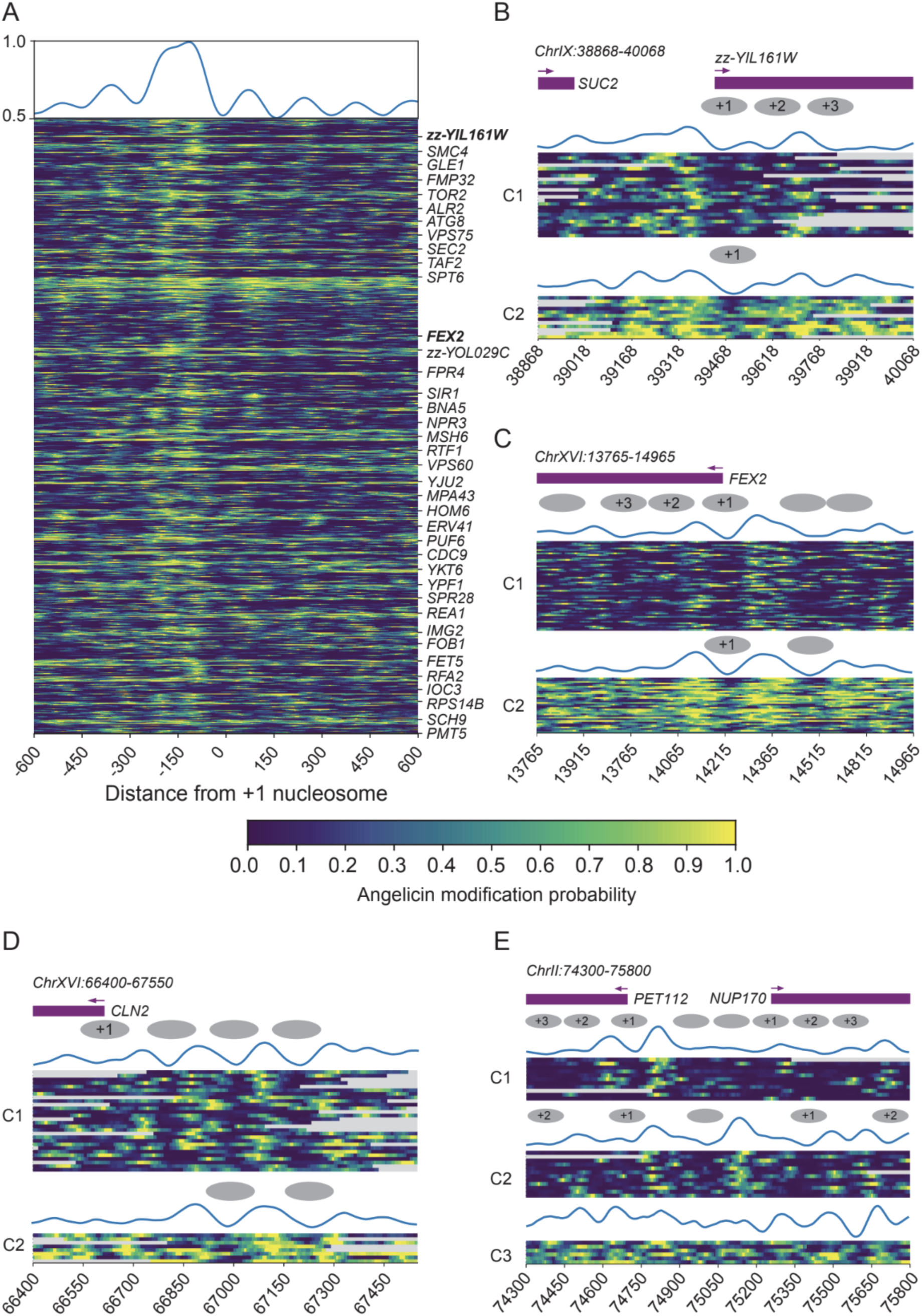
Single-molecule analysis of chromatin structure using SMAdd-seq. Each row is a single DNA molecule covering the locus. The heatmap shows the probability of angelicin modification where yellow is likely modified and blue is unlikely unmodified. The reads are grouped using k-means clustering of the modification scores. The top wiggle shows average modification scores per cluster after min-max normalization. Gray ovals in the schematic represent the predicted positioning of the nucleosomes through visual inspection. (A) Top genes with well-positioned +1 nucleosomes at TSS. Top 38 genes were shared between yeast nuclei replicate 1 and replicate 2 samples and replicate 1 data was used for visualization. Genes were ranked by the Pearson correlation of its modification scores at TSSs with whole genome aggregated modification scores at TSSs (showing replicate 1). (B) zz-YIL161W (ChrIX:38,868-40,068, +) gene promoter region centered at +1 nucleosome dyad. Reads were grouped into 2 clusters. Showing replicate 2. (C) FEX2 (ChrXVI:13765-14965, -) gene promoter region centered at +1 nucleosome dyad. Reads were grouped into 2 clusters. Showing replicate 1. (D) The CLN2 promoter (ChrXVI:66,400-67,550, -). Reads were grouped into 2 clusters. Showing replicate 2. (E) the NUP170 TSS (ChrII:74,300-75,800, +). Reads were grouped into 3 clusters. Showing replicate 1.

When we zoom into specific gene loci, we can cluster the reads aligned to those loci based on the accessibility predictions. *zz-YIL161W* and *FEX2* are two genes with well-structured chromatin based on our analysis (Figure 3B-C). Both genes have 2 clusters of reads with distinct patterns of accessibility that are reminiscent of the structural heterogeneity previously observed for the *PHO5* promoter of yeast by psoralen-EM analysis (Brown et al. 2013). In *zz-YIL161W,* we observe the +1, +2, and +3 nucleosomes in C1, while only the +1 nucleosome is observed in C2, indicating more transcriptionally active chromatin (Figure 3B). In the *FEX2* gene, we observe well-positioned nucleosome arrays in C1 indicating a transcriptionally repressive state while in C2 we only observe the +1 nucleosome and an upstream nucleosome representing a transcriptionally active state of this gene. C2 is also more accessible overall than C1. We can also identify changes in chromatin accessibility in regions other than the TSS. The *CLN2* promoter is a well-studied cell cycle-regulated promoter that has a large nucleosome-depleted region (NDR) upstream of the TATA box (Bai et al. 2010). After k-means clustering of the predicted modification scores at this locus, we observe a cluster (C1) that shows 3 well-positioned nucleosomes in the promoter region and little accessibility downstream (Figure 3D). The second cluster shows only one nucleosome positioned in the promoter and greater accessibility in the gene body, indicating a more transcriptionally active subset of cells regulated by transcription factors binding at the upstream NDR.

We can also identify clustering patterns that are not dependent on overall accessibility. *PET112* and *NUP170* are divergent neighboring genes (Fig 3E). After clustering the reads at this locus, we identify 3 distinct clusters - one with only the *PET112* promoter open (C1), one with both promoters open (C2), and one with higher accessibility across the locus (C3). Compared to C1, both genes in C2 were more accessible, possibly due to greater transcriptional activity. These genes represent examples of a range of chromatin accessibility patterns that can be investigated using SMAdd-seq.

## DISCUSSION

As previously described (Komura et al. 2001), we also find that angelicin can modify thymine bases in a 5’-TA context on single strands of DNA and our work has shown that these strands can be sequenced on nanopores. We also show that angelicin-modified k-mers had a distinct current signal compared to unmodified k-mers. Additionally, we trained a neural network model for estimating the probability of DNA modification by angelicin from nanopore signal measurements. Furthermore, beyond its application in angelicin modification detection, our method can also be used for detecting any kind of modification with a matched positive and negative training data set. Although angelicin modification caused distinct signal shifts in a small fraction of k-mers, our machine learning model was, on average, able to detect open and closed chromatin, different patterns of chromatin accessibility and patterns of intramolecular correlation. These methods allow us to detect both the chromatin accessibility on a genome-wide level and single-molecule level at specific loci. We also show that angelicin can modify DNA and accurately mark chromatin accessibility *in vivo*, without the need for nuclei extraction.

The biggest challenge we have faced with nanopore sequencing of angelicin-modified DNA is the sparseness of data, primarily due to a combination of incomplete modification of accessible modifiable sites and blockage of the pores, presumably due to DNA cross-linking. We were also surprised that we did not detect a bias for 5’-AT, 5’-TG or 5’-GT dinucleotide adducts in our sequencing data. However, both, Esposito et al., (1988) and Komura et al., (2001) also observed a similar dinucleotide bias in their data with psoralen suggesting that the overwhelming preference for modifying 5’-TA that we observed may be a more common property of furocoumarins. Some of the incomplete angelicin modification may also be due to the fact that intercalation results in the random modification of only one strand and not the other in a TA/AT context, because angelicin can only covalently bond with a single thymine base in one strand of the DNA. As a result, we failed to sequence the strand containing the modification half the time with standard nanopore sequencing. This means that even in our positive control, we are not detecting modifications for all modifiable sites. This is a non-trivial problem, especially for the neural network-based model, as machine learning models depend highly on good-quality training data. Other groups have used synthetic DNA with modified bases at known sites to train similar models. However, we could not find any available protocols or companies that could generate synthetic angelicin-modified DNA templates. Future work may utilize the newly developed nanopore duplex sequencing to sequence both strands of DNA (Sanderson et al. 2023a, 2023b), increasing the probability of sequencing the modified k-mer at each modifiable position. However, at the moment, the duplex sequencing method is currently not sufficiently high-throughput enough to generate datasets for network training (Sanderson et al. 2023a, 2023b).

Previous work has suggested that the chemistry of angelicin should not allow for the formation of cross-links (Ashwood-Smith and Grant 1977; Komura et al. 2001), unlike the angelicin analog, psoralen, which mostly forms interstrand cross-links (Brown et al. 2013). However, we observed a small fraction of angelicin-treated DNA containing interstrand cross-links. This result combined with the more rapid decay of flow cell pores on samples with angelicin treatment, leads us to hypothesize that the interstrand cross-links in the DNA cannot pass through the pores, thus clogging them and reducing the throughput of the flow cell. One way to alleviate this issue would be to incubate DNA at elevated temperature and basic pH to break interstrand cross-links. Base treatment has been successfully used before to break DNA cross-links formed by psoralen (Shi, Spielmann, and Hearst 1988). However, the adapter protein required to ratchet the DNA through the nanopore during sequencing will not withstand such harsh treatment thus precluding this option. Conversely, treating the modified DNA with alkali prior to adapter ligation to reverse cross-links would render it single-stranded and since the adapter ligates only to dsDNA, perfectly renaturing long stretches of high-complexity DNA would be challenging.

Other options include modifying angelicin itself as well as generating alternative angular structures of other furocoumarin derivatives to determine how they traverse the nanopore and utilizing this information to synthesize and test alternative small molecules that can be used as probes for visualizing the chromatin landscape (Lampronti et al. 2017; Tupini et al. 2022). Other less damaging furocoumarin derivatives will also enable this method to be extended to alternative long-read sequencing methods like PacBio sequencing since DNA polymerases are currently unable to polymerize through an angelicin-modified template (data not shown).

Despite these challenges, our current protocol allowed us to detect chromatin accessibility both at the genome-wide and single locus level. There are other additional benefits to using this small molecule as opposed to enzymes in probing chromatin structure. Compared to enzyme-based approaches, angelicin modification is significantly cheaper per Gb of sequence generated-approximately $1 for angelicin compared to $100 for the commercial EcoGII methyltransferase based on throughput from (Dennis, Xu, and Clark 2024). Furthermore, angelicin is naturally found only in plant cells and therefore acts as a fully exogenous DNA modification in fungi and animal cells (Mahendra et al. 2020). Other approaches use GpC methyltransferases to label genomes that also have endogenous CpG modification, which results in the exclusion of methylation data in a GCG context due to ambiguity between native methylation and exogenous modification (Lee et al. 2020). Angelicin is also a membrane-permeable molecule, which can facilitate chromatin accessibility probing without isolating nuclei (Komura et al. 2001), which has been previously shown to affect chromatin structure accessibility (X. Wang and Simpson 2001; Prentice and Gurley 1983). Removing the step of nuclei isolation can make accessibility probing more amenable to low-input tissue samples or other single-cell analyses. While the angelicin modification protocol is in need of further optimization, we show that nanopore sequencing of angelicin-modified chromatin is a highly feasible method for probing chromatin structure *in vivo*.

## DATA AVAILABILITY

Raw nanopore signal data are deposited at https://zenodo.org/records/15122707. Basecalled nanopore sequencing data and alignment files are available under BioProject PRJNA1084879.

Data and Codes for regenerating figures are at: https://github.com/baigal628/smaddseq_manuscript. Our computational model NEMO is available at https://github.com/baigal628/NEMO.

## AUTHOR CONTRIBUTIONS

Gali Bai: Data Curation, Formal Analysis, Methodology, Software, Visualization, Writing – original draft, Writing - review & editing

Namrita Dhillon: Methodology, Supervision, Writing – original draft, Writing – review & editing

Colette Felton: Data curation, Formal Analysis, Investigation, Methodology, Project administration, Software, Visualization, Writing – original draft, Writing – review & editing

Brett Meissner: Methodology, Resources, Writing - review & editing

Brandon Saint-John: Conceptualization, Data Curation, Methodology, Investigation, Formal Analysis, Writing - review & editing

Robert Shelansky: Conceptualization, Methodology, Formal Analysis, Writing - review & editing

Eva Hrabeta-Robinson: Conceptualization, Methodology, Funding Acquisition, Project Administration, Writing - review & editing

Elliot Meyerson: Methodology, Software, Writing - review & editing

Babak Hodjat: Conceptualization

Hinrich Boeger: Conceptualization, Funding Acquisition, Supervision, Writing - review & editing

Angela N. Brooks: Conceptualization, Funding Acquisition, Supervision, Project Administration, Writing - review & editing

## FUNDING

This work was supported by the National Institutes of Health [R35GM138122 to A.N.B, T32HG008345 to B.S.-J.]; QB3-UCSC Graduate Innovators Fellowship to G.B.; and the National Science Foundation [#2111763]. Funding for open access charge: National Institutes of Health/R35GM138122 and National Science Foundation/2111763.

## CONFLICT OF INTEREST

E.M. and B.H. are employees of Cognizant. All other authors declare no conflict of interests.

## ACKNOWLEDGEMENTS

We would like to thank Michael Doody and Zhipeng Lu for helpful advice.

## Notes

https://zenodo.org/records/15122707

https://github.com/baigal628/smaddseq_manuscript

https://github.com/baigal628/NEMO

### Summary of Updates

We have changed the method name to Small-Molecule Adduct sequencing (SMAdd-seq). We have improved QC and computational analysis with a comparison to methyltransferase-based long-read methods. Finally, we now demonstrate SMAdd-seq using intact cells, without the need for nuclei isolation.

