## Supplemental Figures for "SMAdd-seq: Probing chromatin accessibility with small molecule DNA intercalation and nanopore sequencing"

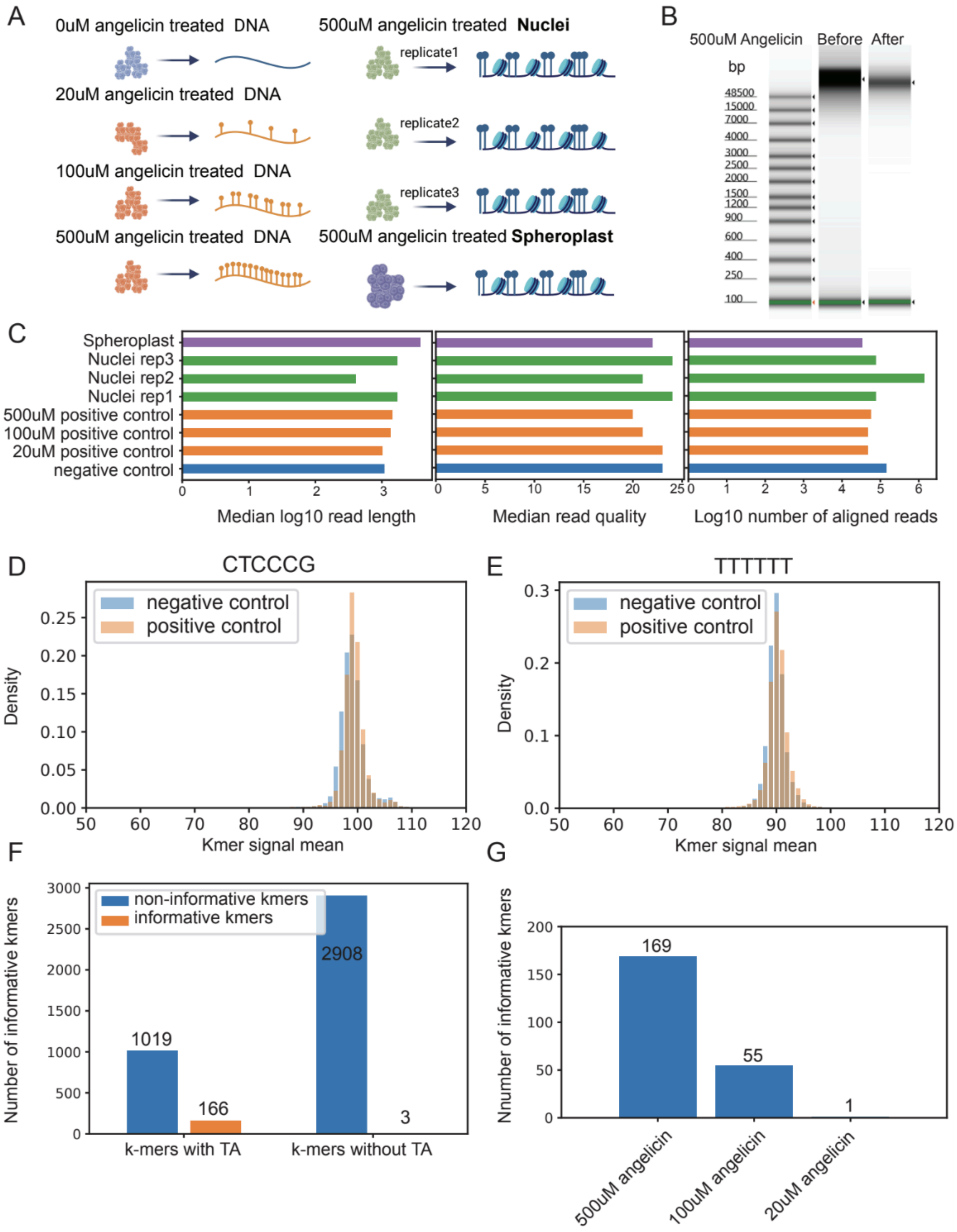

A
0uM angelicin treated DNA
20uM angelicin treated DNA
100uM angelicin treated DNA
500uM angelicin treated DNA
500uM angelicin treated **Nuclei**
replicate1
replicate2
replicate3
500uM angelicin treated **Spheroplast**
B
500uM Angelicin
Before
After
bp
48500
15000
7000
4000
3000
2500
2000
1500
1200
900
600
400
250
100
C
Spheroplast
Nuclei rep3
Nuclei rep2
Nuclei rep1
500uM positive control
100uM positive control
20uM positive control
negative control
Median log10 read length
Median read quality
Log10 number of aligned reads
D
CTCCCG
negative control
positive control
Density
Kmer signal mean
E
TTTTTT
negative control
positive control
Density
Kmer signal mean
F
non-informative kmers
informative kmers
Number of informative kmers
1019
166
2908
3
k-mers with TA
k-mers without TA
G
Nnumber of informative kmers
169
55
1
500uM angelicin
100uM angelicin
20uM angelicin

**Supplementary Figure 1. Data quality control and k-mer analysis**

(A) A summary of yeast sequencing data generated for the SMAdd-seq experiments, which includes a titration experiment on genomic DNA with angelicin concentration from 0uM, 20uM, 100uM to 500uM (left), triplicates of yeast nuclei treated with 500uM angelicin and one replicate of spheroplast treated with 500uM angelicin.

(B) A TapeStation profile of spheroplast DNA treated with 500uM angelicin before (left lane) and after (right lane) library preparation. This sample was used to prepare ONT libraries with the SQK-LSK110 sequencing kit.

(C) Bar plots showing the quality control metrics from nanopore sequencing data. Chromatin rep3 is dropped from downstream analysis due to the lack of genes with well-positioned nucleosomes.

(D & E) Histograms of the nanopore signal currents produced from T containing unmodifiable k-mers in yeast DNA that had been either treated with UV light only (blue: negative control) or with angelicin and UV for (orange: positive control). No shift in signal currents were observed from positive control data.

(F) Bar plots showing number of informative k-mers (orange: has at least one shifted peak in positive control data) and non-informative k-mers (blue: has no shifted peak in positive control data) grouped by whether the kmer contains a modifiable TA dimer.

(G) Bar plots showing increased number of k-mers with detectable modification signal with increased angelicin concentration.

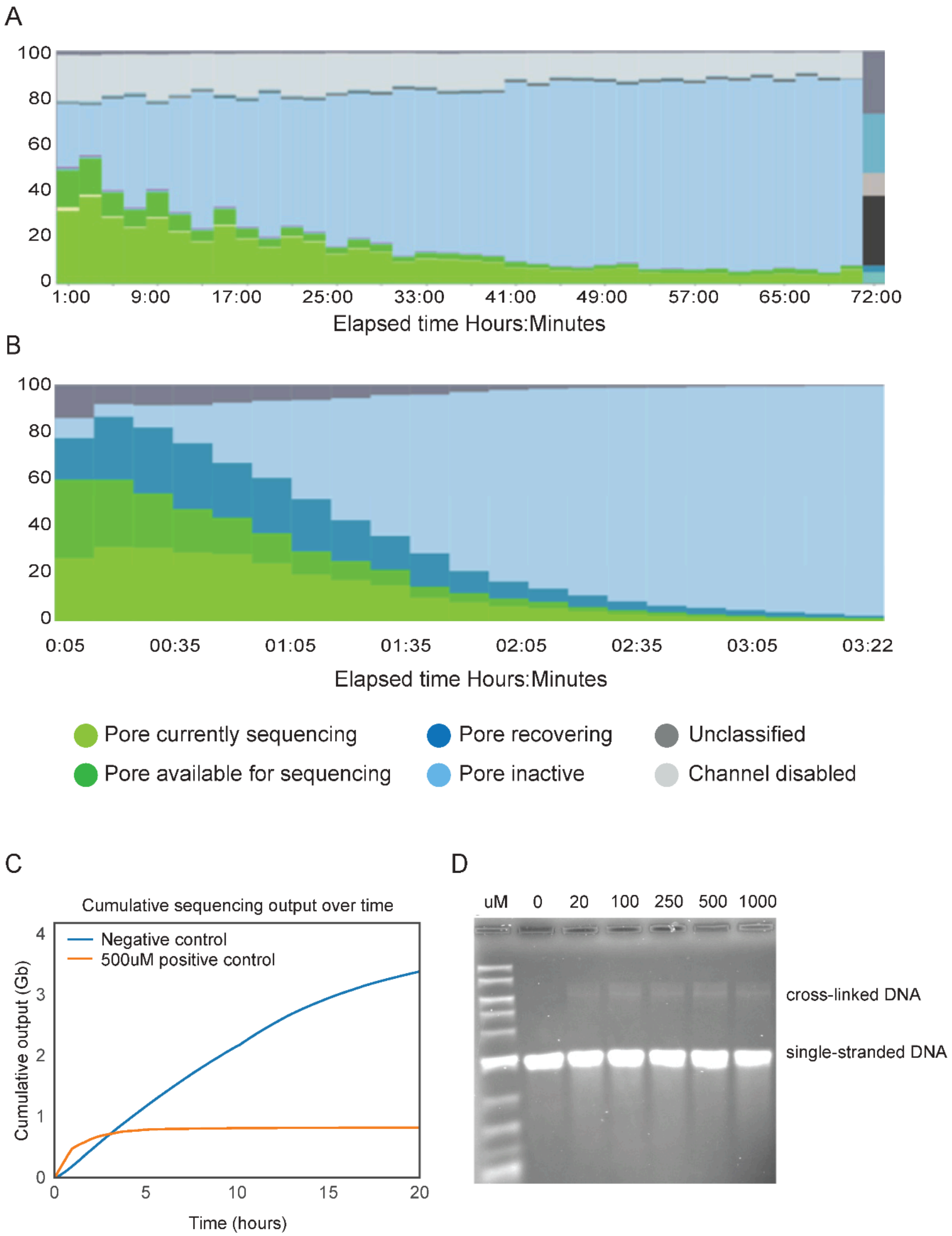

A
100
80
60
40
20
0
1:00
9:00
17:00
25:00
33:00
41:00
49:00
57:00
65:00
72:00
Elapsed time Hours:Minutes
B
100
80
60
40
20
0
0:05
00:35
01:05
01:35
02:05
02:35
03:05
03:22
Elapsed time Hours:Minutes
Pore currently sequencing
Pore available for sequencing
Pore recovering
Pore inactive
Unclassified
Channel disabled
C
Cumulative sequencing output over time
Negative control
500uM positive control
Cumulative output (Gb)
4
3
2
1
0
0
5
10
15
20
Time (hours)
D
uM
0
20
100
250
500
1000
cross-linked DNA
single-stranded DNA

**Supplementary Figure 2. DNA crosslinking due to angelicin modification leads to reduced throughput of flow cells.**

(A & B) The histograms show percent activity of nanopores during a sequencing run with (A) Unmodified DNA, x-axis has a maximum value of 72 hours and (B) DNA modified with 500uM angelicin, x-axis has a maximum value of ~3 hours.

(C) Cumulative sequencing output for a sequencing run of DNA treated with either 0uM or 500uM angelicin. X-axis indicates the time elapsed for sequencing.

(D) Denaturing alkaline agarose gel electrophoresis of linearized plasmid BlueScript (pBS) modified with varying concentrations of angelicin (0uM to 1000uM). A majority of the DNA migrated as single stranded DNA (lower band); however, a small amount of double-stranded DNA (upper band) was observed in lanes with angelicin-modified DNA.

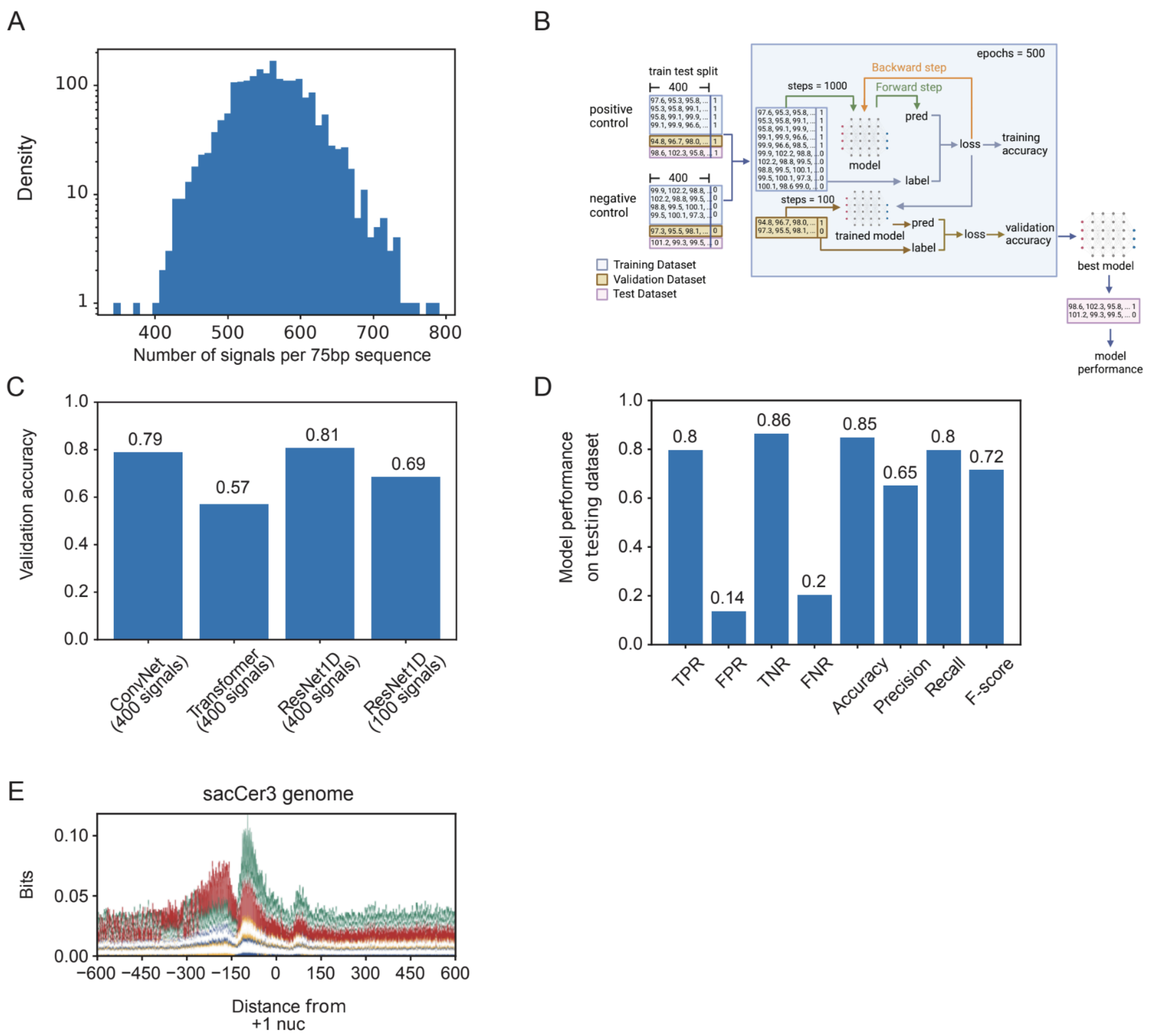


**Supplementary Figure 3. Detailed illustration of deep learning model training and model performance evaluation**

(A) Number of signal scores per 75bp window after eliminating 6-mers with > 10 scores assigned to them (indicates DNA pausing, causes outliers).

(B) A schematic of the model training process. Signal currents from each control sample were split into training (60%), validation (20%), and testing (20%) datasets. After training 500 epochs, the best model was chosen based on the best prediction accuracy on validation dataset. The final model performance is evaluated on the testing dataset.

(C) Bar plot comparing prediction accuracy on validation dataset when using different neural network architectures and different data input sizes.

(D) Bar plot showing final model performance on testing dataset. TPR=true positive rate, FPR=false positive rate, TNR=true negative rate, FNR=false negative rate

(E) Sequence logo showing motif enrichment at transcription start sites in the yeast genome. A depletion of TA was observed at 150 bp upstream +1 nucleosome dyad (A: Green, T: Red, C: Blue, G: Yellow).

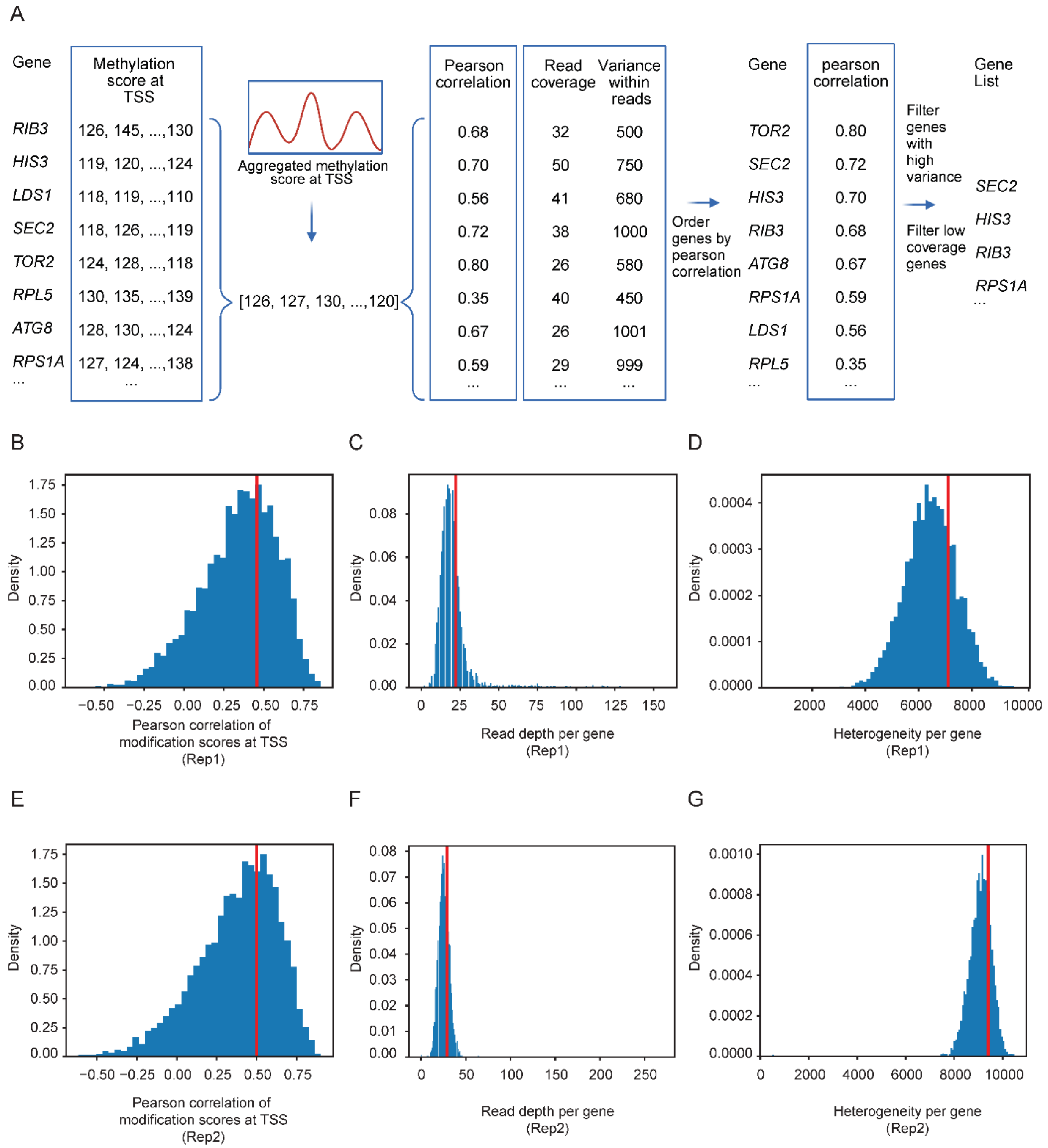


**Supplementary Figure 4. Selection of genes with well-positioned nucleosomes at transcription start sites.**
(A) A schematic illustrating genes with well-positioned nucleosomes were selected based on the Pearson correlations between their angelicin modification scores at TSS and the genome-wide aggregated angelicin modification scores.
(B-G) Histogram showing the distribution of Pearson correlation of modification scores at the promoter of a single gene with the aggregate promoter modification pattern, read depth per gene and heterogeneity between reads per gene across all annotated genes in the Nuclei rep1

sample (B, C & D) and Nuclei rep1 sample (E, F & G). Red lines indicate the 75 percentile cutoffs used for filtering genes.
